# Pain Stickiness in Complex Regional Pain Syndrome: A role for the Nucleus Accumbens

**DOI:** 10.1101/769802

**Authors:** Andrew M. Youssef, Ke Peng, Pearl Kijoo Kim, Alyssa Lebel, Navil F. Sethna, Corey Kronman, David Zurakowski, David Borsook, Laura E. Simons

**Author notes:** David Borsook and Laura E. Simons contributed equally to this manuscript. **Correspondence:** David Borsook, MD PhD, Center for Pain and the Brain, c/o 1 Autumn Street, Boston, MA 02215, e.

## Abstract

Some individuals with chronic pain experience improvement in their pain with treatment, whereas others do not. The neurobiological reason is unclear, but an understanding of brain structure and functional patterns may provide insights into pain’s responsivity to treatment. In this investigation, we used magnetic resonance imaging (MRI) techniques to determine grey matter density alterations on resting functional connectivity (RFC) strengths between pain responders and nonresponders. Brain metrics of pediatric patients at admission to an intensive pain rehabilitative treatment program were evaluated. Pain responders reported significant pain improvement at discharge and/or follow-up whereas nonresponders reported no improvements, increases in pain, or emergence of new pain symptoms. The pain (responder/nonresponder) groups were compared with pain-free healthy controls to examine predictors of pain responder status via brain metrics. Our results show: (1) on admission, pain nonresponders had decreased grey matter density (GMD) within the nucleus accumbens (NAc) and reduced RFC strength between the NAc and the dorsolateral prefrontal cortex vs. responders; (2) Connectivity strength was positively correlated with change in pain intensity from admission to discharge; (3) Compared with pain-free controls, grey matter and RFC differences emerged only among pain nonresponders; and (4) Using a discriminative model, combining GMD and RFC strengths assessed at admission showed the highest prediction estimate (87%) on potential for pain improvement, warranting testing in a de novo sample. Taken together, these results support the idea that treatment responsiveness on pain is underpinned by concurrent brain structure and resting brain activity.

## Introduction

An understanding of how chronic pain may resolve or persist remains unclear. The term “Pain Stickiness” has been used to infer responsivity based on age, sex, genetics, social and psychological factors that may confer responsivity to pain reversal in patients suffering with chronic pain [13]. Here we evaluate responsivity to treatment (responders vs. non-responders) in a cohort of patients with complex regional pain syndrome (CRPS), using brain systems function and structure as a basis that may help understand “stickiness” to resolution of pain. In CRPS patients, differences in brain structure and function have been identified when compared to healthy peers. Studies have revealed grey matter reduction in the nucleus accumbens, insula, prefrontal and cingulate cortices [6; 17; 20]. Among functional brain imaging investigations, mixed results are reported in child and adult populations as compared with pain-free controls. Specifically, in adult populations, wide-spread hypoconnectivity patterns within the default mode network [10] and insula-centered covariation [29], whereas pediatric investigations generally report hyperconnectivity in neural networks [7] and amygdala-centered covariation [53]. Given these, it is possible that these differential patterns may reflect age-related effects, or perhaps a transition of circuitry as a consequence of disease duration. Indeed, there is evidence of a shift from sensory to emotional circuits with disease duration [25]. Taken together, these alterations likely reflect brain indicators of CRPS disease state that may relate to complex processes of the temporal nature (duration of symptoms) of the condition, genetic background and other factors [22] of the disease that integrate emotional and sensory processing, particularly in the context of reward and aversion [12].

As noted above, our group has previously explored structural and functional brain metrics in pediatric CRPS following intensive interdisciplinary pain rehabilitation [7; 17; 53]. One of the observations in the latter reports was the ‘normalization’ of networks and gray matter changes when pretreatment vs. posttreament scans were evaluated. Despite observed changes in brain metrics and converging evidence of improved pain and functioning [26; 33; 56], not all patients respond to treatment with improvements in pain [55]. Despite emerging evidence of brain markers of the transition from acute to chronic pain [4; 25], there is no known data on baseline brain metrics that can distinguish pain responders and nonresponders who undergo pain treatment. Identifying these metrics can further our understanding of drivers of pain treatment resistance and provide potential targets for pharmacotherapy or that may be amenable to early interventions such as motivational interviewing [1] prior to embarking on costly [18] and time intensive pain treatment interventions. Given that reward circuits are considered to be important in pain chronification [4; 25], in pain responsivity to treatment [31], and motivation to change have been identified as a key risk factor for treatment response [55], we hypothesized that patients who are resistant to pain improvement will have significant structurally and functionally altered reward processing regions (viz., NAc) vs. those that are responsive.

## Materials and methods

### Subjects

Twenty-nine patients (8 males, 21 females; mean ±SEM age: 13.5±0.5 years; range 10-20 years) with CRPS of the lower extremity and 29 well-matched healthy controls (8 males, 21 females; mean ±SEM age: 13.7±0.6 years; range 8-21 years) were recruited for the study. All patients were recruited from the Pain Treatment Services (Chronic Pain Clinic) at Boston Children’s Hospital and diagnosed by an experienced neurologist in accordance with a neurological examination and a comprehensive record review. Patients who were then admitted to the Mayo Family Pediatric Pain Rehabilitation Center (PPRC) were enrolled in the study. The day hospital rehabilitation program at the time of enrolment incorporated intensive physical, occupational, and psychological therapy eight hours per day, five days per week for an average length of stay of three to four weeks. A typical treatment day began with a three one-hour blocks of individual physical therapy (PT), occupational therapy (OT), and psychological therapy followed by a two-hour period for studying and lunch. The subsequent two hours consisted of a one-hour of group physical or occupational therapy session and a one-hour of group psychological therapy session. For the final hour of the day, patients participated in either family psychotherapy (twice per week) or parent-observed individual physical or occupational therapy sessions (three times per week). A physician and nurse evaluated patients daily to ensure continued appropriateness for treatment (e.g., continued medical stability) and to address acute or ongoing medical issues. PT, OT, and psychotherapy focused on helping children return to premorbid levels of functioning through progressively engaging in previously avoided activities and taking a self-management approach to pain. Additional details on the program and its outcomes are reported elsewhere [34]. After completing the program patients returned for a follow-up appointment at 1-month post-discharge. At that time, patients were evaluated by a physician, psychologist, occupational therapist, and physical therapist individually, for one hour sessions each. Following these evaluations, the treatment team met and provided the family with feedback regarding current clinical status, goal attainment, and goal progression.

Across time, patients reported to the rehabilitation program nurse their average pain intensity over the past 24-hours on a numerical rating scale (NRS; 0 = no pain, 1-3 = mild pain, 4-6 = moderate pain, 7-10 = severe pain) at admission, discharge and one-month follow up. These numbers were recorded in the patient’s medical record and extracted for this analysis. Individual patients whose pain levels significantly decreased in severity from the time of admission to one-month follow-up (severe to moderate, moderate to mild or mild to none) were categorized as *pain responders*, whereas those whose pain levels either remained unchanged, increased, or developed a new pain problem were classified as *pain nonresponders*.

To assess physical competences from admission to follow-up, patients completed the Functional Disability Inventory (FDI; [63]). To evaluate psychological measures, patients completed the Children’s Depression Inventory (CDI; [57]), Multidimensional Anxiety Scale for Children (MASC; [38]) and the Fear of Pain questionnaire (FOPQ-C; [54]). Patients were excluded from the study if they had any other neurological symptoms, severe medical problems (such as uncontrollable asthma and seizures, cardiac diseases or severe psychiatric disorders), medical implants and/or devices or weighed more than 285 pounds, corresponding to the magnetic resonance imaging (MRI) scanner weight limit. All patients were instructed not to take analgesic medication within at least 4 hours prior to the study scanning session. Healthy controls were recruited through flyers on bulletin boards throughout the local community, online list serves (e.g., craigslist, college job boards) and word of mouth. Informed written consent was obtained for all procedures, which were conducted under the approval of Boston Children’s Hospital Institutional Review Board.

### MRI acquisition

All subjects were positioned supine in a Siemens Magnetom Trio 3 Tesla Magnetic Resonance Imaging (MRI) scanner (Siemens Healthcare Inc., East Walpole, MA, USA), and data acquired using a 12-channel head coil. For image registration, a three-dimension magnetization-prepared rapid gradient-echo (MP-RAGE) sequence was used to acquire a high resolution T1-weighted anatomical image (repetition time = 1410 ms, raw voxel size = 1.0 mm thick, matrix = 256 × 256 voxels). In a subset of these patients (6 responders, 6 nonresponders) and correspondent 12 well-matched controls, a series of 200 gradient echo echo-planar image sets with Blood Oxygen Level Dependent (BOLD) contrast was collected at rest (axial slices = 41, repetition time = 3000 ms, echo time = 35 ms, raw voxel size = 3.75 × 3.75 × 3.50 mm thick, matrix = 64 × 64 voxels). In the remaining patients (13 responders, 4 nonresponders) and correspondent 17 well-matched controls, a 150 gradient echo echo-planer image sets with BOLD contrast was collected (axial slices = 41, repetition time = 2500 ms, echo time = 30 ms, raw voxel size = 3.75 × 3.75 × 3.50 mm thick, matrix = 64 × 64 voxels) was collected. Here, it is imperative to note that these data were collected over eight years and acquiring two differential sequences, although not ideal for this investigation, was employed to aquire a faster repetition time to correspond with the current trend of sequence acquisition. This would also allow enchance the resolution of other functional imaging techniques such as measuring fluctuations in low frequency amplitude, whereby the bandwidth is correspondent to the repetition time.

### MRI analysis

#### Grey matter density analysis

Using SPM12 [19], image preprocessing and grey matter density maps were created with the computational anatomy toolbox (dbm.neuro.uni-jena.de/cat). First, to optimize skull-stripping and spatial normalization for our pediatric population, we created a customized tissue probability map and template using the Template-O-Matic toolbox [65]. The toolbox allows for the construction of high-quality templates from structural data derived from four-hundred healthy subjects aged five to eighteen years [65]. In two of the fifty-eight subjects who exceeded the age-range (i.e. age twenty), we allocated a value of eighteen. All T1-weighted images were then resliced (0.5mm^3^) and segmented into grey matter (GM), white matter (WM) and cerebrospinal fluid (CSF) probability maps. Here, the segmentation approach was based on the Adaptive Maximum A-Posterior (AMAP) technique to model local variations and parameters as slowly varying spatial functions [44] and the Partial Volume Estimation (PVE) method whereby estimating the fraction of each tissue type within each voxel (GM-WM and GM-CSF) to yield more accurate segmentation [59]. Furthermore, to enhance the quality of the T1-weighted images and further improve the segmentation, we used two denoising methods, combining the Spatially Adaptive Non-Local Means (SANLM) filter [37] with the classical Markov Random Field (MRF) method [44].

The segmented images were then registered to the previously created pediatric tissue probability map using affine transformation (i.e. linear, preserving proportions), followed by a non-linear deformation in Montreal Institute Neurological (MNI-152) space. The non-linear deformation parameters were calculated by using the high-dimensional Diffeomorphic Anatomical Registration Through Exponentiated Lie Algebra (DARTEL) algorithm [2; 3] with the previously created study-specific pediatric template. To correct for expansion (or contraction) during the spatial transformation, the normalized images were then “modulated” by multiplying each voxel by the Jacobian determinant (i.e. linear and non-linear components) derived from the spatial normalization procedure. Finally, the normalized, modulated grey matter images were then smoothed using an isotropic 8 mm full-width-half-maximum (FWHM) Gaussian filter and these images used for subsequent analysis.

Significant differences in grey matter density between responders and nonresponders were determined within a general linear model using an independent two-sample t-test. To account for the differences in brain size between subjects, we calculated the total intracranial volume for each subject and included these values as a nuisance variable. In addition, age and gender were factored out by modelling them as nuisance variables. Following an *a priori* primary threshold of p<0.001, we applied a family-wise error rate (FWE) cluster-level extent threshold to correct for multiple comparisons. We chose this stringent cluster-defined primary threshold based on the simulations of Woo and colleagues [66] to optimize the control of false positives and to improve overall inferences about specificity. Significant linear relationships between grey matter density values and disease duration, pain intensity (admission) and the percentage change in pain intensity (from admission to discharge) were determined using Pearson’s partial correlation analysis, while controlling for age and total intracranial volume influences (p<0.05).

#### Functional connectivity analysis

Using SPM12 [19], image preprocessing and seed-based functional connectivity maps were created with the functional connectivity toolbox [64]. The first five volumes were discarded to allow for T1-equilibration effects and all images subsequently realigned (rigid body translation and rotation) to the first volume as the initial motion correction procedure. Next, to correct for slice-dependent time shifts during image acquisition, we applied slice-time correction for all the functional data. The slice-time corrected images were then coregistered to the same subjects’ T1-weighted anatomical image and the anatomical image spatially normalized to Montreal Neurological Institute Space (MNI-152). The deformation field acquired during affine and non-linear transforms of the T1-weighted anatomical image were then applied to the functional images (i.e. indirect normalization). Finally, the normalized functional images were spatially smoothed with an isotropic 8mm full-width-half-maximum (FWHM) Gaussian kernel. Images were linearly detrended to remove global signal intensity changes and a temporal high pass-filter with a 100 second time constant applied.

To account for nonspecific variance, eighteen physiological and motion-related related factors were included as nuisance variables. These included the first three principle components of the time course derived from separate regions of matter and cerebral spinal fluid [8] and the six body translation and rotation parameters from the realignment procedure. The first temporal derivative of the movement parameters was included to account for temporal shifts in the signal. In recent years, a whole-body of literature has established that in-scanner head movement can have substantial influences on resting state functional connectivity [43; 60], which is particularly problematic in pediatric populations [50]. Here, particular care was taken with examining movement; for all subjects, we set an *a prior* criterion of less than 3mm cumulative displacement and 3 degrees of angular motion. One patient and two healthy controls exceeded this criterion and were therefore excluded from the functional connectivity analyses. Additionally, the remaining subjects’ head motion were examined, by computing the maximum absolute translational movement and angular rotation [60] and scalar frame wise displacement [43] derived from the six rigid body parameters. Here, the purpose of these measures is to index head movement, not to precisely model it or set additional exclusionary criteria.

Using the normalized smoothed images, regions where significant grey matter density values between responders and nonresponders were used as “seeds” in the subsequent resting state analysis. The seed-based functional connectivity map was generated by computing the correlation coefficient between each voxels time series with the mean time course of the voxels within the seed region. The individual correlation coefficient maps were then converted to z-maps using Fisher’s r-to-z transformation and these images used for group-level statistical comparisons. Significant differences between responders and nonresponders were then determined using a two-way analysis of variance (ANOVA) under the framework of a general linear model. We chose this model to ensure that any significant group differences observed were not dependent on differential repetition times (i.e. no interaction effect) during image acquisition. Here, we included between-subjects’ factor (group: responders; nonresponders) and within-subjects’ factor (repetition time: slow; fast). Gender and age were also factored out by modelling them as nuisance variables. Significant main and interaction effects were determined using an *a priori* primary threshold of p<0.001 with a family-wise error rate cluster-level extent threshold applied to correct for multiple comparisons. Post-hoc two-sample t-tests were then conducted between the groups to determine the directionality of main effect of group (p<0.05). Significant linear relationships between resting seed-based functional connectivity values and disease duration, pain intensity (admission) and the percentage change in pain intensity (from admission to discharge) were determined using Pearson’s partial correlation analysis, while controlling for age (p<0.05).

#### Imaging metrics in healthy controls

To examine grey matter density and subsequent seed-based functional connectivity values in well-matched healthy controls, we overlaid the results derived from the between-patients contrasts described above. Here, we extracted grey matter density values and resting seed-based functional connectivity from overlapping clusters. Significant differences between patients (responders and nonresponders) and well-matched healthy controls were then determined using independent two-sample t-tests (p<0.05).

#### Discriminative model and predictive estimate analysis

We conducted discriminant analyses to assess potential model performance using the extracted imaging metrics in a more intuitive and quantitative way. We hypothesized that imaging metrics would surpass age, pain complexity, and physical or psychological functions in predicting a patient’s pain responsivity status. These factors were included in the comparisons as they were likely to also have predictive influence on patient pain responsivity, as shown in our previous work [55]. Here, we used a support vector machine-based linear discriminative model with different features to segregate pain responders and nonresponders. For imaging metrics, we used grey matter density and resting connectivity strengths metrics derived from the voxel-based morphometry and seed-based functional connectivity contrasts (see above). The performance of the discriminative model in providing predictive estimates on the responsivity of new patients was evaluated through a Monte-Carlo resampling approach [67]. Specifically, the acquired imaging metrics of the patients were first randomly partitioned into a training set (containing 80% of the total data points) and a test set (containing the remaining 20% of data points). The discriminator was trained with the training dataset and the parameters of the linear discriminator were selected using a leave-one-out cross-validation approach. We then applied the trained discriminator to the test dataset to determine *predictions* on whether a patient in the test set would have pain improvement. For each group of features, the random partitioning, training and predicting process was repeated two hundred times. In each repetition, the results were compared with their outcome behaviour to calculate the number of patients on which correct and incorrect predictions were obtained using the following definitions: *True Positive* (TP), the discriminator correctly marked a patient whose pain improved as a responder; *True Negative* (TN), the discriminator correctly marked a patient who did not have pain improvement as a nonresponder; *False Positive* (FP), the discriminator incorrectly marked a patient who did not have pain improvement as a responder; *False Negative* (FN), the discriminator incorrectly marked a patient who experienced pain improvement as a nonresponder.

In this study, we compared the results using three different groups of imaging metrics as discriminative features, including (1) both grey matter density and functional connectivity values, (2) grey matter density only and (3) functional connectivity only. Patient baseline characteristics that were included in the comparisons are: (4) age, (5) pain intensity at admission, (6) pain duration, (7) physical measure at admission (FDI score), (8) psychological measure at admission (CDI, MASC and FOP scores). For each group, the average prediction accuracy estimate, sensitivity and specificity (defined below) across repetitions were reported:

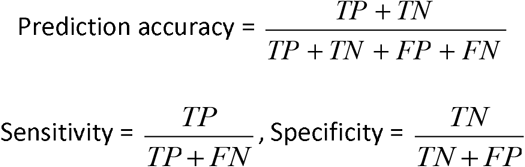

## Results

Individual responder and nonresponder subject characteristics, imaging modality inclusion, pain duration and pain intensity are shown in **Table 1** and between-group comparisons in **Fig. 1A**.

**Table 1:** Clinical characteristics of patients. Significant differences in pain characteristics between pain responders and nonresponders are colored grey, determined by independent two-sample t-test (p<0.05). F = female; M = male; B = bilateral; U = unilateral; L = left; R = right. *Removed from grey matter density analysis due to poor image aquisition **Removed from seed-based connectivity analysis due to excessive head motion.

| Patient | Age | Gender | Pain duration (months) | Site | Reported pain levels (0-10) at: |  |  | Complicating factors (following discharge) | Classification |
| --- | --- | --- | --- | --- | --- | --- | --- | --- | --- |
|  |  |  |  |  | Admission | Discharge | Follow Up |  |  |
| 1 | 17 | F | 3 | R | 7 | 5 | not reported | knee injury pain spread to lower leg and foot | nonresponder |
| 2 | 10 | F | 19 | R | 7 | 6.5 | 6 | right hip pain | nonresponder |
| 3 | 11 | M | 8 | L | 3 | 1.5 | 0 |  | responder |
| 4 | 15 | F | 5 | B | 7 | 6.5 | 8 | memory loss, headaches, paralysis | nonresponder |
| 5 | 15 | M | 12 | R | 7 | 1.5 | 1.5 |  | responder |
| 6 | 11 | F | 7 | L | 6 | 0 | 0 |  | responder |
| 7 | 16 | F | 48 | L | 2.5 | 2 | 0 |  | responder |
| 8 | 17 | F | 19 | L | 6 | 2 | 1 |  | responder |
| 9 | 14 | F | 3 | L | 9.5 | 7.5 | 8.5 | depression and suicidal ideation | nonresponder |
| 10** | 13 | M | 6 | B | 4 | 0 | 0 |  | responder |
| 11 | 17 | F | 85 | R | 6 | 5.5 | 6 | right hip pain | nonresponder |
| 12 | 13 | F | 13 | L | 5.5 | 7.5 | 1 | chronic migraines, back pain, vertigo, loss of consciousness, conversion disorder | nonresponder |
| 13 | 12 | F | 8 | B | 8 | 6.5 | 0 |  | responder |
| 14 | 10 | M | 5 | R | 7 | 8 | 0 |  | responder |
| 15 | 15 | M | 11 | L | 4 | 3 | 5.5 |  | nonresponder |
| 16 | 14 | F | 3 | R | 3.5 | 1.5 | 0 |  | responder |
| 17 | 15 | F | 9 | L | 8 | 6 | 5 |  | responder |
| 18 | 15 | F | 10 | R | 7 | 5 | 0 |  | responder |
| 19 | 12 | F | 12 | L | 9 | 8 | 3 |  | responder |
| 20 | 11 | M | 3 | L | 6 | 7 | 3 |  | responder |
| 21 | 11 | M | 3 | R | 7 | 5 | 0 |  | responder |
| 22 | 12 | F | 8 | L | 4 | 3 | 1 |  | responder |
| 23 | 11 | F | 5 | L | 6 | 2 | 0 |  | responder |
| 24 | 20 | F | 47 | B | 8 | 8 | 9 | development of pain at site of walking aid | nonresponder |
| 25 | 12 | F | 8 | R | 4 | 6 | 6 |  | nonresponder |
| 26 | 17 | F | 6 | L | 5.5 | 0 | 0 |  | responder |
| 27 | 11 | F | 3 | B | 3 | 3 | 0 |  | responder |
| 28* | 13 | M | 34 | L | 4 | 1 | 1 |  | responder |
| 29 | 11 | F | 3 | R | 3 | 4 | 8 |  | nonresponder |
| R<br>mean ±<br>SEM | 13.0±<br>0.5 |  | 11.0±2.7 |  | 5.61±0.44 | 3.32±0.63 | 0.82±0.32 |  |  |
| NR<br>mean ±<br>SEM | 14.4±<br>1.0 |  | 19.5±8.4 |  | 6.00±0.63 | 5.95±0.5 | 6.44±0.8 |  |  |

**Figure 1:**
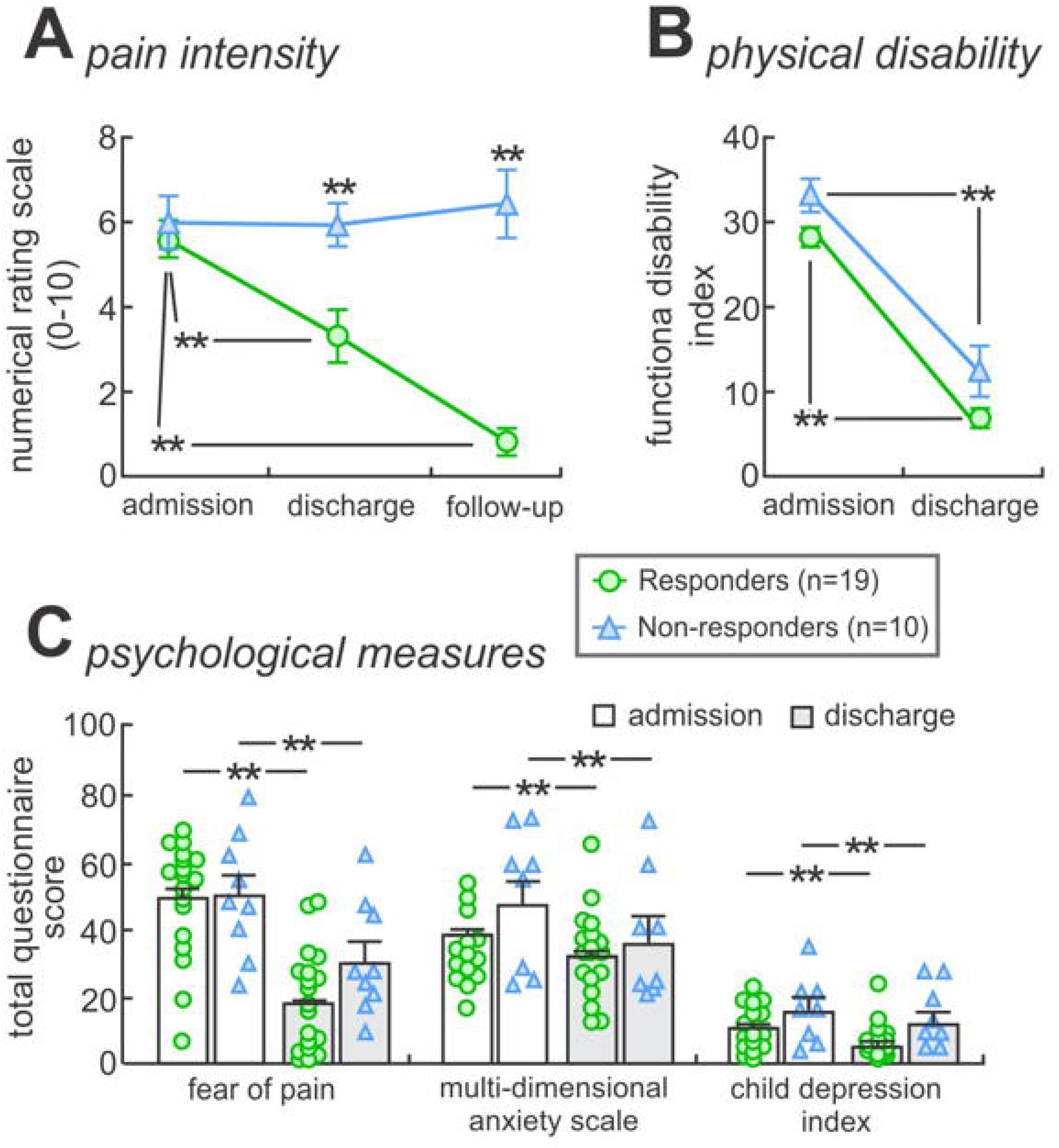
Pain intensity, physical function and psychological measures. **A:** Patients reported their pain intensity on a numerical rating scale (0 = no pain, 1-3 = mild pain, 4-6 = moderate pain, 7-10 = severe pain) at three time points (admission, discharge and one-month follow up). Individual patients whose pain levels significantly decreased in severity from the time of admission to follow-up (severe to moderate, moderate to mild or mild to none) were categorized as pain responders (n=19), whereas those whose pain levels either remained unchanged or increased were classified as pain nonresponders (n=10). Significant between and within-group differences were determined using independent two-sample t-test and paired-t test (**p<0.01). **B:** Total functional disability inventory in responders and nonresponders at admission and discharge. Significant were determined using repeated measures anova (**p<0.01). **C:** Psychological measures (fear of pain, multi-dimensional anxiety scale and child depression inventory) at admission and discharge. Significant differences were determined using repeated measures anova (**p<0.01). Note that no significant group by time interaction was observed.

### Psychometrics

#### Pain Scores

Of the twenty-nine patients, 19 reported a significant reduction in their subjective pain from the time of admission to follow-up (*responders*; mean [±SEM] reduction in pain intensity: − 87.7±4%, paired t-test: p<0.0001), whereas 10 did not (*nonresponders*; mean [±SEM] change in pain intensity: 16.5±9%, paired t-test: p=0.55). The significance equated to at least a 3-point drop in their NRS score. There were no group differences in age (*responders*; 13.0±0.5 years, *nonresponders*; 14.4±1.0; p=0.17), pain duration (*responders*; 11.0±3 months, *nonresponders*; 19.5±8; p=0.23) or pain intensity at admission (*responders*; 5.6±0.4, *nonresponders*; 6.0.5±0.6; p=0.61). In contrast, significant between group differences were observed in pain intensity at discharge (*responders;* 3.32±0.6, *nonresponders;* 5.95.5±0.5; p=0.009) and follow-up (*responders;* 0.82±0.3, *nonresponders;* 6.44±0.8; p<0.0001).

#### Psychological Questionnaires

In addition, to assess potential influences of physical and psychological measures, patients responded to questionnaires at admission through to follow-up. Although most patients completed physical and psychological measures at admission (91%) and discharge (93%), questionnaire scores collected at follow-up were limited to a minor subset (45%). To allow for reliable measures and sufficient power, we therefore confined our statistical comparisons of total physical and psychological measures to admission and discharge using repeated measures ANOVA. Significant decreases in total questionnaire scores (mean [±SEM]) were observed for functional disability inventory (*responders*, admission: 28.8±0.6, discharge: 6.9±0.3; *nonresponders*, admission: 33.2±1.9, discharge: 13.4±3.3, f(20,1)=97.0, p<0.01, δ=0.83), fear of pain (*responders*: 49.7±1.0, 18.8±0.8; *nonresponders:* 51.3±6.4, 31.3±6.0, f(24,1)=37.5, p<0.01, δ=0.61), multidimensional anxiety scale (*responders*, admission: 38.8±1.0, discharge: 32.2±0.7; *nonresponders*, admission: 47.7±7.9, discharge: 37.4±7.2, f(22,1)=8.51, p<0.01, δ=0.28), and child depression inventory (*responders*, admission: 11.4±0.4, discharge: 5.00±0.3; *nonresponders*, admission: 17.1±3.8, discharge: 12.4±3.4, f(22,1)=14.7.0, p<0.01, δ=0.40) (**Fig. 1B, C**). The time by group interaction was non-significant for all outcomes (functional disability p=0.62; fear of pain p=0.20, anxiety p=0.53, depression p=0.56). Thus, pain responders and nonresponders did not differ on self-report measures of emotion and physical function. ra

### Brain Imaging Metrics in Patients

#### Grey matter density

Within the nucleus accumbens [30], pain responders displayed significantly greater grey matter density as compared with nonresponders (mean [±SEM] probability*volume; responders: 0.559±0.009, nonresponders: 0.0478±0.011; p=0.00001) (**Table 2, Fig. 2A**). There were no other significant grey matter density changes in responders as compared with nonresponders. Furthermore, there was no significant difference in the total intracranial volume between these groups (mean [±SEM] cm^3^; responders: 1571±34.3, nonresponders: 1479±23.0; p=0.08). Within the NAc, no significant correlations were observed in grey matter density with pain intensity at admission (r=-0.21, p=0.40), disease duration (r=-0.10, p=0.61) and the percentage change in pain intensity from admission to discharge (r=-0.31, p=0.11) (**Fig. 2B, C**).

**Table 2:**
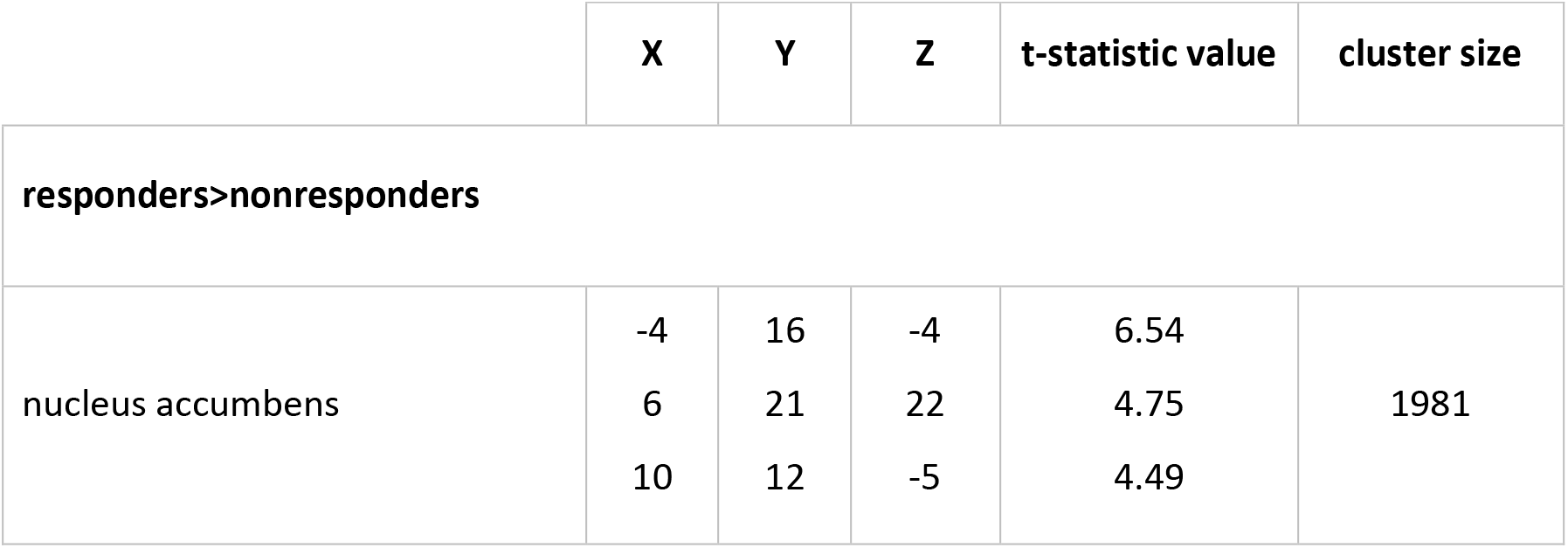
Regions in which grey matter density was significantly different in responders as compared with nonresponders. Locations are in Montreal Neurological Institute space. Note that results were derived from cluster-extent thresholding (family-wise error rate) to correct for multiple comparisons, and therefore, low spatial specificity. Here, peak coordinates are presented as a guide and not recommended for subsequent analyses, instead the cluster should be taken as a whole, that is, predominately encompassing the nucleus accumbens.

**Figure 2:**
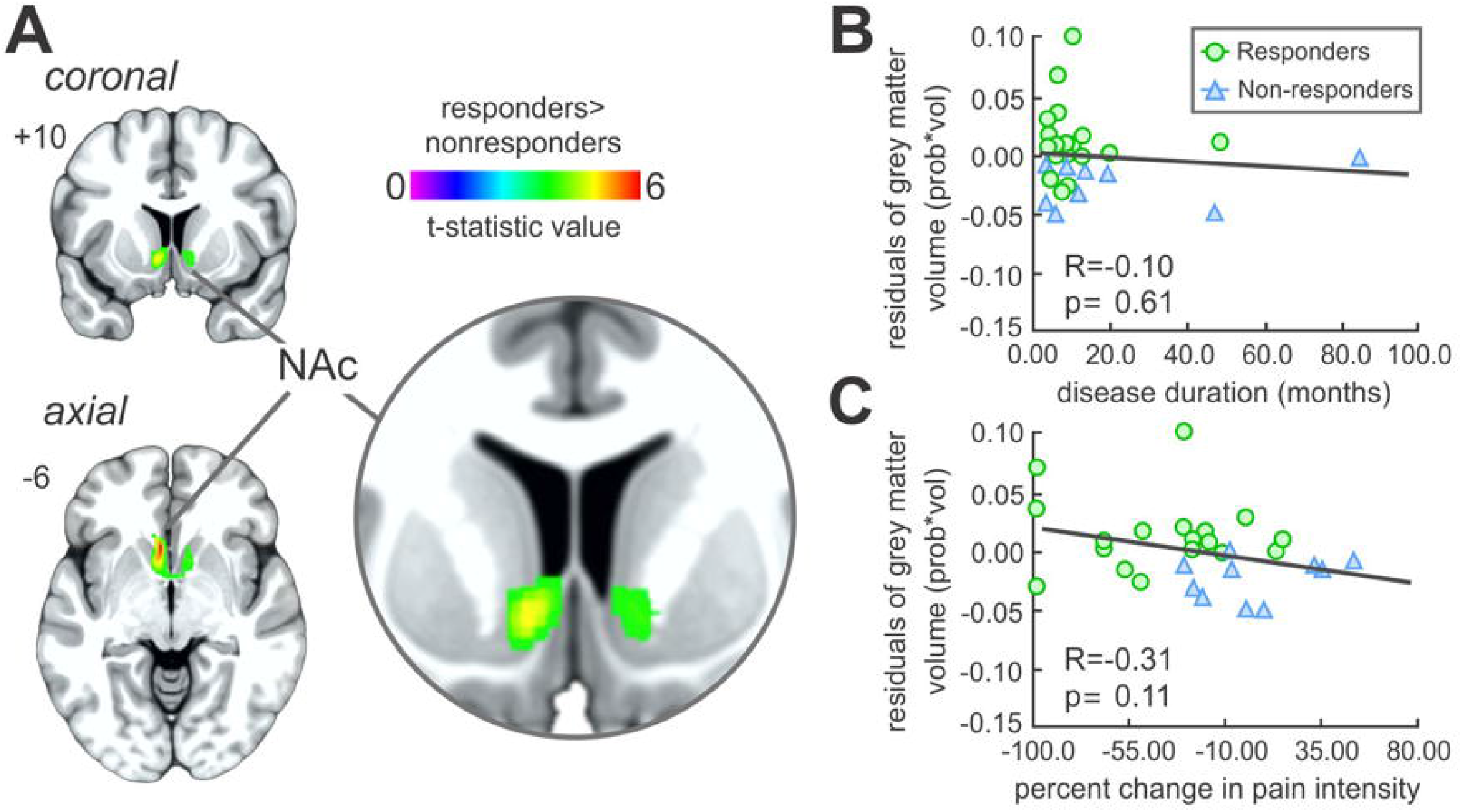
Whole brain grey matter density analysis. **A:** Regional grey matter density reduced within the nucleus accumbens (rainbow colour bar) in pain nonresponders as compared with responders, overlaid onto a coronal and axial T1-weighted anatomical image set. Slice locations are located on the top left of each image and are in Montreal Neurological Institute space. **B:** Partial correlation between regional grey matter density values (controlling for age and total intracranial volume) within the nucleus accumbens and disease duration. **C:** Partial correlation between regional grey matter density values (controlling for age and total intracranial volume) within the nucleus accumbens and percentage change in pain intensity from admission to discharge. Note that no significant partial correlations were observed with clinically reported pain characteristics. NAc, nucleus accumbens.

#### Resting Functional Connectivity

The strength of NAc resting functional connectivity revealed a significant group-type main effect within the right dorsolateral prefrontal cortex (dIPFC) (F=28.78, p<0.0001) (**Table 3, Fig. 3A**). There was no significant main effect of repetition time in responders as compared with nonresponders. Furthermore, there was no significant group-by-repetition-time interaction, demonstrating that group differences observed were not dependent on differential volume acquisition time. Decomposing the main group effect revealed increased NAc-dlPFC functional connectivity in responders as compared with nonresponders (mean [±SEM] parameter estimate values; responders: 0.11±0.04, nonresponders: −0.18±0.07; two-sample t-test, p=0.0001) (**Fig. 3B, C**). Extracting the parameter estimate values of the NAc-dlPFC connectivity strengths revealed no significant correlations with pain intensity at admission (r=0.17, p=0.39) and disease duration (r=0.003, p=0.987) (**Fig. 3D**). In contrast, we observed a negative correlation between NAc-dlPFC connectivity strength and the percentage change in pain intensity from admission to discharge (r=-0.48, p=0.009) (**Fig. 3E**).

**Table 3:**
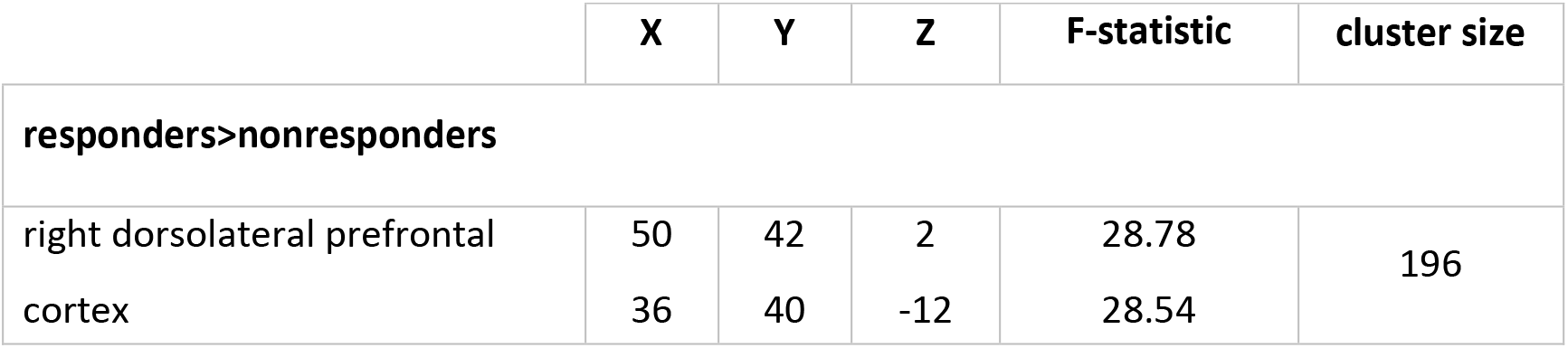
Regions in which resting functional connectivity strengths were significantly different in responders as compared with nonresponders. Locations are in Montreal Neurological Institute space. The nucleus accumbens “seed” was derived from the grey matter density analysis between responders and nonresponders.

**Figure 3:**
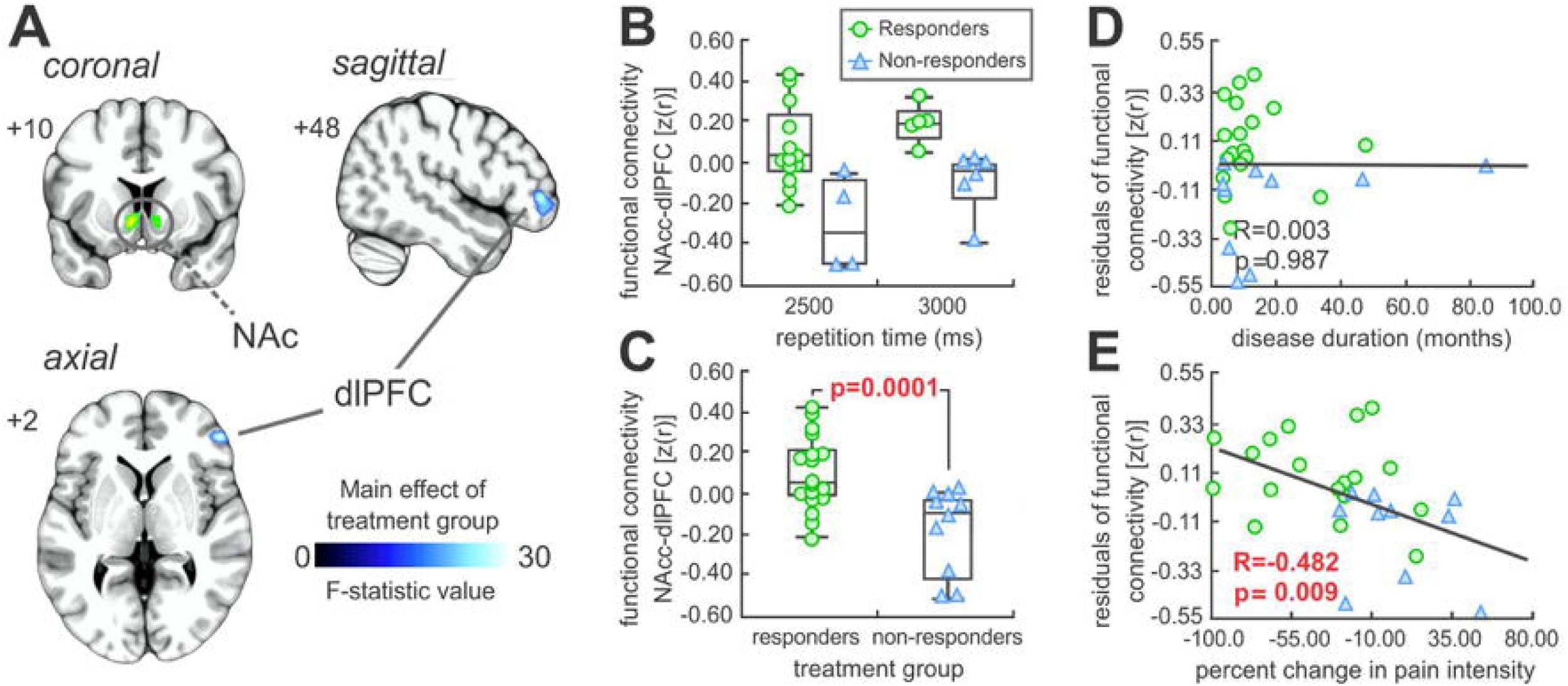
Nucleus accumbens seed-based functional connectivity analysis. **A:** Significant main effect between responders and nonresponders (cool scale bar) in resting functional connectivity strength within the dorsolateral prefrontal cortex, overlaid onto a coronal, axial and sagittal T1-weighted anatomical image set. Slice locations are located on the top left of each image and are in Montreal Neurological Institute space. **B:** Plots of functional connectivity values, note that there was no significant main effect of repetition time or interaction with group-type in responders as compared with nonresponders. **C:** Plots of functional connectivity values for responders and nonresponders. Post-hoc significant differences between the patient groups were determined using an independent two-samples t-test, with the p-value above the plots. **D:** Partial correlation between functional connectivity values (controlling for age) and disease duration. **E:** Partial correlation between functional connectivity values (controlling for age) within and percentage change in pain intensity from admission to discharge. NAc, nucleus accumbens; dIPFC, dorsolateral prefrontal cortex.

#### Movement Artefact

Movement did not significantly impact our results as there were no significant movement-related differences between responders and nonresponders. Here, we investigated (1) mean frame-wise displacement relative to the previous time-point (mean [±SEM] mm; responders: 0.21±0.04, nonresponders: 0.13±0.01; two-sample t-test, p=0.13) [43] and (2) the maximum absolute displacement of the head relative to the origin time-point in translation (mean [±SEM] mm; responders: 1.83±0.35, nonresponders: 1.63±0.29; two-sample t-test, p=0.73) and rotation (mean [±SEM] degrees; responders: 1.84±0.32, nonresponders: 1.80±0.42; two-sample t-test, p=0.95) [60].

### Brain imaging Metrics in Healthy Controls

#### Grey Matter Density

Grey matter density and seed-based functional connectivity values were extracted in healthy controls from clusters derived from the previous responders and nonresponder analyses described above. Here, healthy controls displayed increased grey matter density values within the NAc as compared with nonresponders (mean [±SEM] probability*volume; controls: 0.548±0.012; two sample t-test, p=0.0018), whereas no difference was observed when compared with responders (p=0.53) (**Fig. 4A**). Similarly, healthy controls were associated with significantly greater NAc-dlPFC functional connectivity as compared with non-nonresponders (mean [±SEM] parameter estimate values; controls: 0.028±0.03; two-sample t-test, p=0.0029), with no difference observed when compared with responders (p=0.12) (**Fig. 4B**). Finally, it is pertinent to note that no significant movement-related differences were observed between controls (frame-wise displacement: 0.15±0.02 mm; absolute displacement: 1.50±0.17 mm; absolute rotation: 1.32±0.43 degrees) as compared with responders (two-sample t-test: p=0.12, p=0.40, p=0.16, respectively) and nonresponders (p=0.40, p=0.75, p=0.43, respectively).

**Figure 4:**
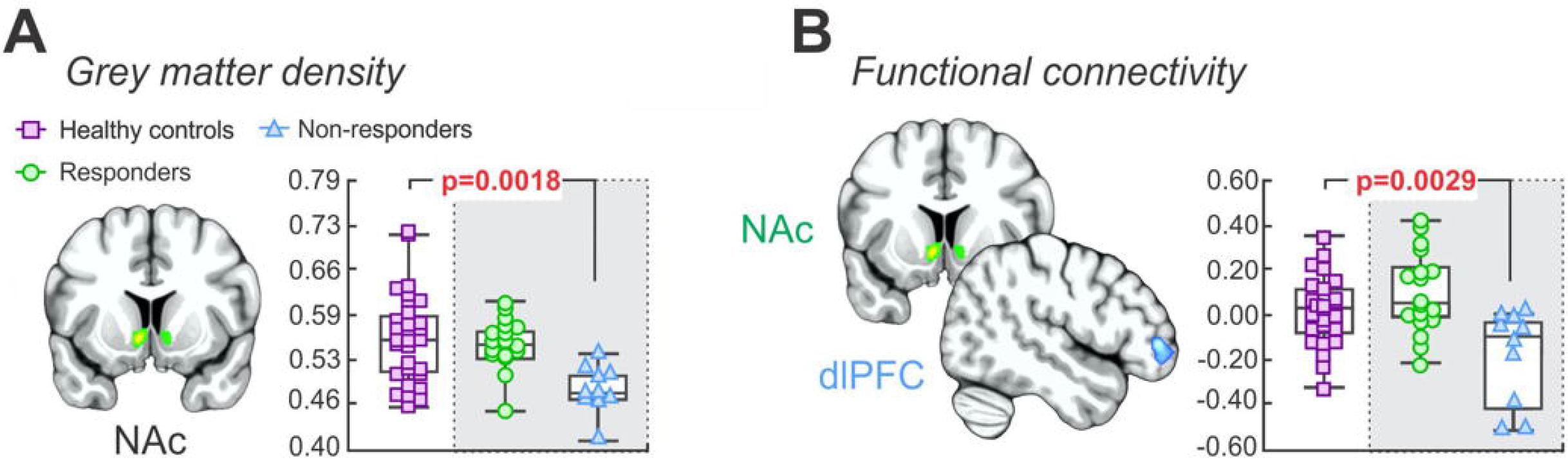
Nucleus accumbens grey matter density and seed-based functional connectivity in healthy controls. **A:** Plots of nucleus accumbens grey matter density values derived from the significant cluster between responders and nonresponders. Significant differences between pain-free controls as compared with responders and nonresponders were determined using independent two-samples t-test (p<0.05), with the p-value above the plots. Note that healthy controls displayed increased grey matter density values as compared with nonresponders, whereas no difference was observed when compared with responders. **B:** Plots of resting seed-based functional connectivity from overlapping clusters derived from the significant cluster between responders and nonresponders. Similarly, healthy controls were associated with significantly greater functional connectivity as compared with non-nonresponders with no difference observed when compared with responders. NAc, nucleus accumbens, dIPFC, dorsolateral prefrontal cortex.

### Discriminative and Predictive Estimate Analysis

The NAc grey matter density and NAc-dlPFC functional connectivity values were used to determine the cross-validated linear discriminative model (responders, n=17; nonresponders, n=10) (**Fig. 5A**). In the predictive estimate analysis, these twenty-seven data points were randomly divided into a training set (responders, n=13; nonresponders, n=8) and a test set (responders, n=4; nonresponders, n=2). Using both the grey matter density and functional connectivity values as discriminative features; we obtained a higher accuracy of (mean [±SEM]) 87±0.8%, sensitivity of 89±1% and specificity of 84±0.2% than those using only grey matter density (accuracy = 84±0.9%, sensitivity = 87±1%, specificity = 78±2%) or functional connectivity (accuracy = 76±1%, sensitivity = 78±2%, specificity = 71±3%) (Mann Whitney U-test p-value < 0.001). The relatively higher sensitivity and lower specificity reveal that the discriminative model with image metrics exhibited better performance in identifying pain responders than nonresponders patients (**Fig. 5B**). Moreover, these obtained overall prediction accuracy using image metrics were significantly higher than those obtained using patient baseline characteristics (p-values < 0.001). **Fig. 5C** shows the Receiver Operating Characteristic [42] curves and their corresponding 95% confidence intervals of the discriminators estimated from the predictive estimate analysis. The area under curves (AUCs) of the ROC curves were compared with a non-parametric approach [15], revealing significant larger AUCs values of the ROC curve using both the grey matter density and functional connectivity data than using only one of the metrics (p-value < 0.001), and using patient baseline characterisitics (p-values < 0.001). Internal validation based on a random selection of 80% training data and 20% of test data using Monte-Carlo resampling with 200 repetitions indicated that the combination of grey matter density and functional connectivity provided the most robust prediction and best internal validity in differentiating pain responders from nonresponders (AUC = 0.930, 95% CI: 0.908-0.948) [40].

**Figure 5:**
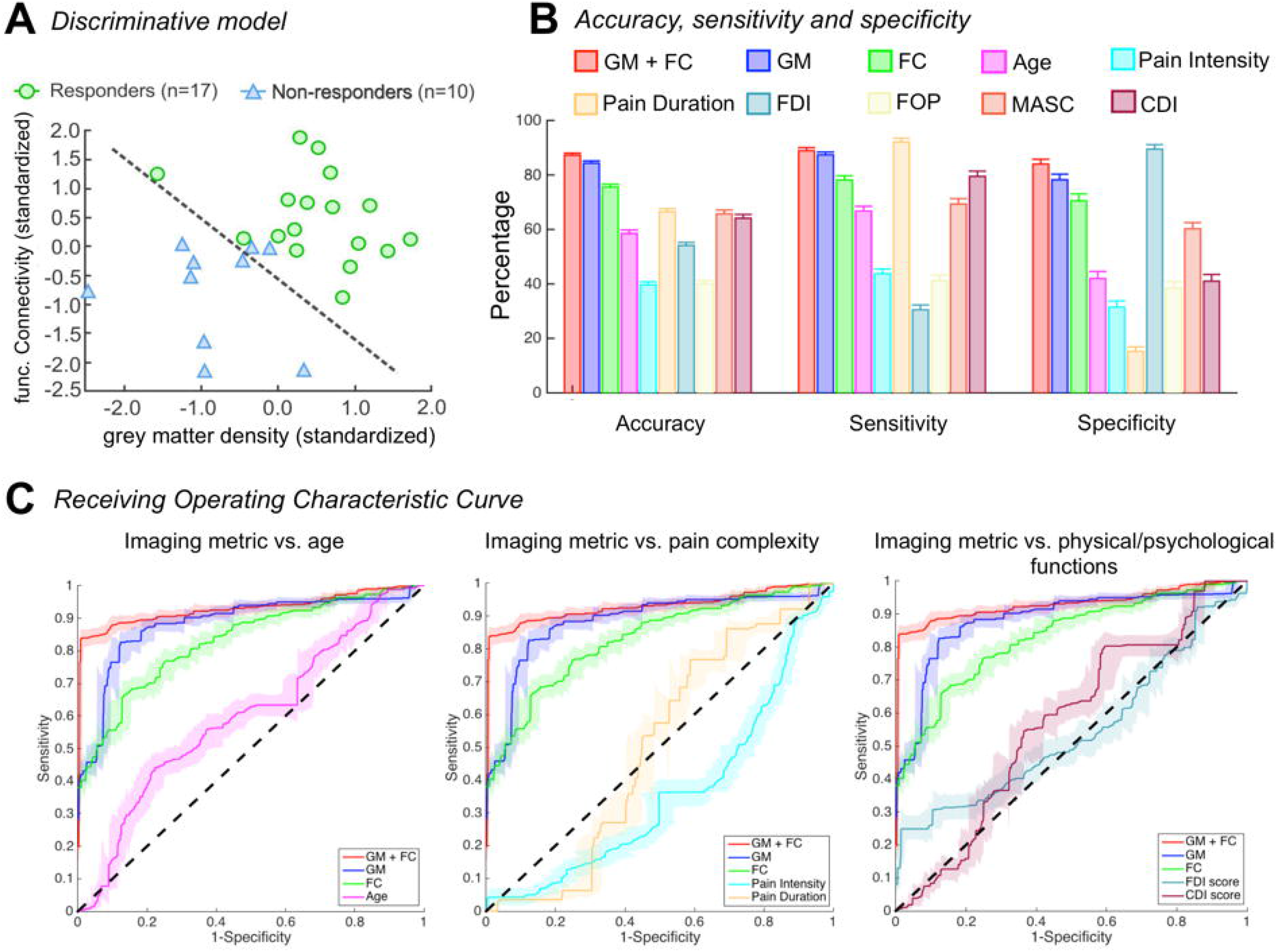
Discriminative model and predictive estimate analysis. **A:** Depiction of discriminative model using both the NAc grey matter density and NAc-dlPFC functional connectivity of the twenty-seven patients where the model was cross-validated using a leave-one-out approach. **B:** The accuracy, sensitivity and specificity of predicting pain responder patients of different discriminators, using imaging metrics including both grey matter density and functional connectivity, only the grey matter density, only the functional connectivity, as well as patient baseline characteristics such as patient age, pain complexity (pain intensity and duration) at admission, physical and psychonological functions at admission as features. These values were obtained by averaging the results of the two hundred permutation tests. Error bars represent standard error of the mean. **C:** Comparisons of the estimated Receiver Operating Characteristic (ROC) Curves using the three imaging metric discriminators in the predictive analysis vs. baseline characteristic discriminators. NAc, nucleus accumbens; dIPFC, dorsolateral prefrontal cortex, GM, grey matter density; FC, functional connectivity; FDI, functional disability inventory; FOP, fear of pain; MASC, multidimentional anxiety scale for children; CDI, children’s depression inventory. For image clarity, only the ROC curve of CDI scores was shown in the comparison of psychological factors.

### Summary of Results

Our results in a sample of pediatric patients with CRPS suggest that structural and functional alterations can predetermine a patients propensity for pain improvement in treatment. Specifically, at admission, pain responders were associated with increased grey matter density within the nucleus accumbens as compared with nonresponders. No associations between grey matter density and disease duration, pain intensity or change in pain intensity from admission to discharge were observed. Similarly, responder patients had greater resting functional connectivity strengths between the NAc and the dorsolateral prefrontal cortex (dIPFC), and this finding was not associated with disease duration or pain intensity at admission. In contrast with NAc grey matter density, these resting functional connectivity strengths were positively correlated with the change in pain intensity from admission to discharge. In pain nonresponders, patients had reduced grey matter density and functional connectivity metrics as compared with pain-free controls. Our predictive estimate analysis showed that the magnitudes of NAc grey matter density and functional connectivity strengths could potentially predict whether an individual would have pain improvement in response to treatment with an accuracy of 87%. Taken together, these results support the idea that pain improvement in treatment is underpinned by concurrent brain structure and resting brain activity.

## Discussion

To the best of our knowledge, no study to date has evaluated brain structural or functional differences between pain responders and nonresponders in CRPS. Our data reveal two major findings: (1) That NAc gray matter volume is different in pain responders vs. nonresponders prior to intensive treatment; and (2) NAc grey matter density and resting functional connectivity strengths with the dIPFC can potentially predict, with 87% accuracy, pain responsiveness in patients with CRPS.

### NAc Morphometry in Pain Response in CRPS

In this investigation we found, compared with controls, reduced NAc grey matter density in pain nonresponders, whereas no differences were observed in responders. Given this, it is possible that a lack of consistency across investigations may reflect, in part, differences in patient’s hedonic tone, anhedonic state and/or diminished motivational drive that are often comorbid with chronic pain syndromes [52]. Of course, we do not neglect influences of age, symptomology, pain distribution, intensity and duration across these investigations on brain morphometry [39]. Instead, our study provides a unique perspective; given similar pain characteristics, we report a reduction in grey matter density within the NAc in pain nonresponders as compared with responders. Furthermore, we report analogous findings exclusively in nonresponder patients when compared with pain-free controls. Our data provide one of the reasons that may account for differences across investigations, and we suggest that separating patients into two groups on the basis of the presence or absence of responsiveness is a more robust method of detecting underlying brain changes.

We have previously examined structural brain changes in pediatric patients CRPS following an intensive multidisciplinary pain treatment service [17]. Here, despite reductions in pain intensity, functional disability, fear of pain, anxiety and depression, no NAc changes were observed from admission to discharge [17], possibly reflecting a lack of association with these measures. This is consistent with the findings of this study, that is, despite NAc structural alterations, no differences were observed in these psychophysical measures between responders and nonresponders. Furthermore, we did not find any association between NAc grey matter density and pain intensity at admission. Additionally, since we did not observe an association between the NAc and disease duration, our results may reflect predisposing factors that contribute to a patient’s resilience to pain chronification rather than a consequence of the disease. Alternatively, a lack of NAc treatment-induced differences may reflect ongoing processes that result in delayed NAc structural alterations outside the study time frame [17]. Indeed, Baliki and colleagues [4] compared recovering and persistent subacute back pain, where concurrent morphometric differences within the NAc and patient’s negative affective state were not observed until one year thereafter. Taken together, these data support the idea that the NAc does not seem to encode pain sensory information, but rather may reflect differences between patient’s pain affective state.

### NAc and Functional Connectivity with dlPFC

Using the NAc morphological difference as a seed, pain nonresponders were associated with decreased resting functional connectivity strengths exclusively within the dlPFC, as compared with responders. Over recent years, human brain imaging and lesion investigations have highlighted a diverse range of roles for the dlPFC; driving appropriate behavioural goals by rendering sensorimotor information [41], operating working memory to resist distracting stimuli [49], processing conflicting stimuli efficiently [36], and, in the context of pain, placebo analgesia [30; 62], modulatory processes [35; 68], tolerance [23], thresholds [11], and perceived intensity and unpleasantness [35]. In contrast with the NAc, we did not report dlPFC morphometric alterations between responders and nonresponders. This is consistent with adult populations, that is, no study to date has reported altered dlPFC grey matter density changes as compared with pain-free controls [5; 6; 20; 61]. However, it is pertinent to note that despite a lack of structural alterations, Barad and colleagues [6] reported an association between grey matter density with disease duration and perceived pain intensity. If this is the case, since we report no differences in these pain characteristics between responders and nonresponders, this may, in part, explain a lack of structural difference observed in this study.

In contrast with adult populations, we have previously reported reduced dIPFC cortical thickness as compared with pain-free controls in pediatric populations [17]. Furthermore, following treatment, cortical thickening within the dIPFC and enhanced functional connectivity with the (pain modulatory) periaqueductal grey matter was reported [17]. Notwithstanding these contributions, it is pertinent to note that Gennatas and colleagues [21] recently proposed that metrics of grey matter density may be more sensitive to age-related changes than measures of cortical thickness. Of course, in this study we compared patient responders and nonresponders and therefore, whether or not alterations in cortical thickness differ within patients remains unclear. In addition to structural metrics, alterations in resting functional connectivity strengths in several brain networks in CRPS patients have also been reported [7]. For example, signal covariation within frontoparietal and central executive networks that include, but not limited to, the dIPFC in pediatric CRPS as compared within pain-free controls and these related to treatment effects [7]. Furthermore, amydala-related functional connectivity with the dIPFC was altered in pediatric CRPS and were associated with pain-related fear [53]. In this study, we report alterations in resting functional connectivity strengths between the NAc and the dIPFC, and instead may reflect alterative characteristics. Given the diverse range of functions between the NAc and the dIPFC (see above), the covariation between these brain sites in our study may reflect may reflect behavioural dominance of pain that is dependent upon motivational and reward circuitry. In our study, we did not find any association between NAc-dlPFC functional connectivity and disease duration or pain intensity. Indeed, more recently, Reddan and Wager [45] proposed that the NAc and dIPFC does not appear to be associated with stimulus intensity. In contrast, our results indicate that these connectivity strengths may predispose an individuals pain responsiveness. This idea is discussed further below.

### Predictive Metrics of Pain Stickiness

We found that resting connectivity strengths between the NAc and dlPFC were associated with the change in pain intensity following discharge. Therefore, it may be that while NAc morphometry predisposes the individual, modulatory influences over the NAc by the dlPFC appears to be a primary determinant. Indeed, we found that concurrent structural and functional metrics were associated with greater prediction accuracy, sensitivity and discrimination than structural and functional metrics in isolation. Remarkably, we were able to predict with approximately 87% accuracy whether an individual will express a robust pain improvement based on the NAc grey matter density and resting functional connectivity strengths with the dlPFC. The prediction accuracy using brain metrics were also seen to be higher than those using age, pain intensity at baseline or patient physical and psychrological characteristics. Our previous study based on a large group of patients (n = 253) suggested the predictive influence of several baseline characteristics (e.g. pain problem complexity, older adolescence age and cognitive-affective factors) [55]. However, results from this work showed that brain metric measures might have a better performance in the prediction of individual responsivity, especially when only limited prior information is available. Here, it is possible that the interaction of the dlPFC with the NAc may thus reflect the active manipulation of the behavioural dominance of pain dependent upon motivational and reward context. It is possible that the dlPFC may protect the maintenance of reward and aversive behavioural goals by rendering working memory operations resistant to distracting stimuli [49] and efficient performance in the presence of conflicting stimuli.

## Limitations

We note the following limitations of this study: (1) <u>Gender</u>: Given that our cohort was predominantly female, we did not have the statistical power to explore gender-related differences. As noted above, the condition affects girls more frequently than boys and this represents a ‘normative’ distribution. Although, we cannot exclude the possibility that some of the observed changes may be related to hormonal/menstrual changes in our subjects; (2) <u>Technical</u>: For our functional connectivity analysis, we combined scanning parameters with differential repetition times. Though we cannot completely exclude these influences on our results, no significant interaction effect was observed between our groups and repetition time, and therefore we are confident that our results were not primarily determined by this influence; <u>Cohort Size</u>: The relative number of subjects per group is relatively low, and in part a reflection of the inherent difficulty of performing pediatric studies, notably in children with chronic pain. This also applies to the discriminative model and the predictive estimate analysis conducted on this relatively small sample size; (4) <u>Discriminative model</u>: these techniques generally require a large number of data points to establish a reliable discriminative performance. Here, it is pertinent to note that the reported 87% of accuracy, 89% of sensitivity and 84% specificity using both imaging metrics were obtained based on predictions of individual responsitivity. The current approach does not account for the effect of previous probabilities on the predictability of our features [47]. For example, the base rate of responders in the CRPS patient group was previously estimated to be around 75% [27; 55]. Applying this base rate via Bayes’ theorem to the reported sensitivity (89%) and specificity (84%) leads to a positive predictive value (PPV) of 94.3% and a negative predicitive value (NPV) of 71.8% in identifying a responder in the CRPS patient population. Moreover, taking the incidence rate of CPRS (reported to be around 0.025%; [14]) into the model results in a PPV of 0.019% and a NPV of 0.05% in identifying a CRPS responder patient in the general population (assuming healthy subjects have comparable brain measures as CRPS responders, see Fig. 4). Therefore, due to the small sample size in our predictive analysis and the lack of estimates of PPVs/NPVs across different populations, the diagnosis utility of our proposed imaging markers should be strictly limited. Future work with a larger and also balanced dataset is required to validate the feasibility of using imaging metrics to predict patient pain improvement; (5) <u>Treatment engagement</u>: daily assessment of treatment engagement was not measured in this pain rehabilitation program, but may have served as a phenotypic marker of pain responder status. Future work should include clinician assessment of treatment engagement (e.g., Pittsburgh Rehabilitation Participation Scale; [32]) as a means of predicting earlier in the course of care a potential correlate to pain improvement; (6) No control group: given the context of intensive pain rehabilitative treatment, it is not possible to have a control group and determine if patients would simply improve without treatment, but our prior work demonstrates that it is not likely [56].

## Conclusions

Prior studies have indicated change in gray matter in responses to treatment including our own on structural and functional changes in CRPS patients [7; 17] and in migraine [28] functional connectivity predicting placebo response in chronic pain [58] or to treatment responsivity in headache [46] or hip replacement [24; 48]. Gray matter changes noted in chronic pain have been reviewed elsewhere [51]. Since differences in pain improvement or conversely pain chronification may relate to interactions with reward centers such as the nucleus accumbens and the dorsolateral prefrontal cortex (DLPFC), mechanisms for responsivity may relate to a number of factors that include innate brain systems (i.e., genetic), or behavioral such as reward learning [9; 16]. Our result indicates that the NAc may be a brain region that evaluates disparate neural processing and is sensitive to defining pain stickiness. Our findings may guide future approaches for modulating related circuitry in individuals prior to treatment interventions. Large scale data sets should allow for predictive measures in patients at the initiation of therapy to define a rational treatment paradigm, but also to provide a more extensive/aggressive paradigm in those patients defined to be at risk for pain “getting stuck”.

## Acknowledgments

The authors declare no compenting financial interests. Research reported in this study was supported by NIH grants (R01NS065051; R01HD083133 to DB and R01HD083270 to LS) and the MayDay Fund, New York (DB). The authors thank Natalia Lopez for helping edit the manuscript.

## References

[1] Alperstein D, Sharpe L. The Efficacy of Motivational Interviewing in Adults With Chronic Pain: A Meta Analysis and Systematic Review. The journal of pain: official journal of the American Pain Society 2016;17(4):393–403.

[2] Ashburner J. A fast diffeomorphic image registration algorithm. NeuroImage 2007;38(1):95–113.

[3] Ashburner J, Friston KJ. Computing average shaped tissue probability templates. NeuroImage 2009;45(2):333–341.

[4] Baliki MN, Petre B, Torbey S, Herrmann KM, Huang L, Schnitzer TJ, Fields HL, Apkarian AV. Corticostriatal functional connectivity predicts transition to chronic back pain. Nature neuroscience 2012;15(8):1117–1119.

[5] Baliki MN, Schnitzer TJ, Bauer WR, Apkarian AV. Brain Morphological Signatures for Chronic Pain. PloS one 2011;6(10):e26010.

[6] Barad MJ, Ueno T, Younger J, Chatterjee N, Mackey S. Complex Regional Pain Syndrome Is Associated With Structural Abnormalities in Pain-Related Regions of the Human Brain. The Journal of Pain 2014;15(2):197–203.

[7] Becerra L, Sava S, Simons LE, Drosos AM, Sethna N, Berde C, Lebel AA, Borsook D. Intrinsic brain networks normalize with treatment in pediatric complex regional pain syndrome. NeuroImage: Clinical 2014;6:347–369.

[8] Behzadi Y, Restom K, Liau J, Liu TT. A component based noise correction method (CompCor) for BOLD and perfusion based fMRI. NeuroImage 2007;37(1):90–101.

[9] Berger SE, Vachon-Presseau É, Abdullah TB, Baria AT, Schnitzer TJ, Apkarian AV. Hippocampal morphology mediates biased memories of chronic pain. Neuroimage 2018;166:86–98.

[10] Bolwerk A, Seifert F, Maihofner C. Altered resting-state functional connectivity in complex regional pain syndrome. J Pain 2013;14(10):1107–1115.e1108.

[11] Borckardt JJ, Smith AR, Reeves ST, Weinstein M, Kozel FA, Nahas Z, Shelley N, Branham RK, Thomas KJ, George MS. Fifteen minutes of left prefrontal repetitive transcranial magnetic stimulation acutely increases thermal pain thresholds in healthy adults. Pain Research & Management: The Journal of the Canadian Pain Society 2007;12(4):287–290.

[12] Borsook D, Linnman C, Faria V, Strassman AM, Becerra L, Elman I. Reward deficiency and anti-reward in pain chronification. Neuroscience & Biobehavioral Reviews 2016;68:282–297.

[13] Borsook D, Youssef AM, Simons L, Elman I, Eccleston C. When pain gets stuck: the evolution of pain chronification and treatment resistance. Pain 2018;159(12):2421–2436.

[14] de Mos M, de Bruijn AG, Huygen FJ, Dieleman JP, Stricker BH, Sturkenboom MC. The incidence of complex regional pain syndrome: a population-based study. Pain 2007;129(1–2):12–20.

[15] DeLong ER, DeLong DM, Clarke-Pearson DL. Comparing the areas under two or more correlated receiver operating characteristic curves: a nonparametric approach. Biometrics 1988;44(3):837–845.

[16] Elman I, Borsook D. Common Brain Mechanisms of Chronic Pain and Addiction. Neuron 2016;89(1):11–36.

[17] Erpelding N, Simons L, Lebel A, Serrano P, Pielech M, Prabhu S, Becerra L, Borsook D. Rapid Treatment-Induced Brain Changes in Pediatric CRPS. Brain structure & function 2016;221(2):1095–1111.

[18] Evans JR, Benore E, Banez GA. The Cost-Effectiveness of Intensive Interdisciplinary Pediatric Chronic Pain Rehabilitation. Journal of pediatric psychology 2016;41(8):849–856.

[19] Friston KJ, Holmes AP, Worsley KJ, Poline JP, Frith CD, Frackowiak RSJ. Statistical parametric maps in functional imaging: A general linear approach. Human Brain Mapping 1994;2(4):189–210.

[20] Geha PY, Baliki MN, Harden RN, Bauer WR, Parrish TB, Apkarian AV. The brain in chronic CRPS pain: abnormal gray-white matter interactions in emotional and autonomic regions. Neuron 2008;60(4):570–581.

[21] Gennatas ED, Avants BB, Wolf DH, Satterthwaite TD, Ruparel K, Ciric R, Hakonarson H, Gur RE, Gur RC. Age-Related Effects and Sex Differences in Gray Matter Density, Volume, Mass, and Cortical Thickness from Childhood to Young Adulthood. The Journal of Neuroscience 2017;37(20):5065–5073.

[22] George E, Elman I, Becerra L, Berg S, Borsook D. Pain in an era of armed conflicts: Prevention and treatment for warfighters and civilian casualties. Prog Neurobiol 2016;141:25–44.

[23] Graff-Guerrero A, Gonzalez-Olvera J, Fresan A, Gomez-Martin D, Mendez-Nunez JC, Pellicer F. Repetitive transcranial magnetic stimulation of dorsolateral prefrontal cortex increases tolerance to human experimental pain. Brain research Cognitive brain research 2005;25(1):153–160.

[24] Gwilym SE, Filippini N, Douaud G, Carr AJ, Tracey I. Thalamic atrophy associated with painful osteoarthritis of the hip is reversible after arthroplasty: a longitudinal voxel-based morphometric study. Arthritis and rheumatism 2010;62(10):2930–2940.

[25] Hashmi JA, Baliki MN, Huang L, Baria AT, Torbey S, Hermann KM, Schnitzer TJ, Apkarian AV. Shape shifting pain: chronification of back pain shifts brain representation from nociceptive to emotional circuits. Brain 2013;136(9):2751–2768.

[26] Hechler T, Wager J, Zernikow B. Chronic pain treatment in children and adolescents: less is good, more is sometimes better. BMC pediatrics 2014;14:262.

[27] Hirschfeld G, Hechler T, Dobe M, Wager J, von Lützau P, Blankenburg M, Kosfelder J, Zernikow B. Maintaining lasting improvements: one-year follow-up of children with severe chronic pain undergoing multimodal inpatient treatment. Journal of pediatric psychology 2013;38(2):224–236.

[28] Hubbard CS, Becerra L, Smith JH, DeLange JM, Smith RM, Black DF, Welker KM, Burstein R, Cutrer FM, Borsook D. Brain Changes in Responders vs. Non-Responders in Chronic Migraine: Markers of Disease Reversal. Frontiers in human neuroscience 2016;10:497.

[29] Kim JH, Choi SH, Jang JH, Lee DH, Lee KJ, Lee WJ, Moon JY, Kim YC, Kang DH. Impaired insula functional connectivity associated with persistent pain perception in patients with complex regional pain syndrome. PloS one 2017;12(7):e0180479.

[30] Krummenacher P, Candia V, Folkers G, Schedlowski M, Schönbächler G. Prefrontal cortex modulates placebo analgesia. PAIN^®^ 2010;148(3):368–374.

[31] Lebel A, Becerra L, Wallin D, Moulton EA, Morris S, Pendse G, Jasciewicz J, Stein M, Aiello-Lammens M, Grant E, Berde C, Borsook D. fMRI reveals distinct CNS processing during symptomatic and recovered complex regional pain syndrome in children. Brain 2008;131(Pt 7):1854–1879.

[32] Lenze EJ, Munin MC, Quear T, Dew MA, Rogers JC, Begley AE, Reynolds CF, 3rd. The Pittsburgh Rehabilitation Participation Scale: reliability and validity of a clinician-rated measure of participation in acute rehabilitation. Archives of physical medicine and rehabilitation 2004;85(3):380–384.

[33] Logan DE, Carpino EA, Chiang G, Condon M, Firn E, Gaughan VJ, Hogan M, Leslie DS, Olson K, Sager S, Sethna N, Simons LE, Zurakowski D, Berde CB. A Day-hospital Approach to Treatment of Pediatric Complex Regional Pain Syndrome: Initial Functional Outcomes. The Clinical journal of pain 2012.

[34] Logan DE, Carpino EA, Chiang G, Condon M, Firn E, Gaughan VJ, Hogan M, Leslie DS, Olson K, Sager S, Sethna N, Simons LE, Zurakowski D, Berde CB. A day-hospital approach to treatment of pediatric complex regional pain syndrome: initial functional outcomes. The Clinical journal of pain 2012;28(9):766–774.

[35] Lorenz J, Minoshima S, Casey KL. Keeping pain out of mind: the role of the dorsolateral prefrontal cortex in pain modulation. Brain 2003;126(Pt 5):1079–1091.

[36] MacDonald AW, 3rd, Cohen JD, Stenger VA, Carter CS. Dissociating the role of the dorsolateral prefrontal and anterior cingulate cortex in cognitive control. Science (New York, NY) 2000;288(5472):1835–1838.

[37] Manjon JV, Coupe P, Marti-Bonmati L, Collins DL, Robles M. Adaptive non-local means denoising of MR images with spatially varying noise levels. Journal of magnetic resonance imaging: JMRI 2010;31(1):192–203.

[38] March JS, Parker JD, Sullivan K, Stallings P, Conners CK. The Multidimensional Anxiety Scale for Children (MASC): factor structure, reliability, and validity. Journal of the American Academy of Child and Adolescent Psychiatry 1997;36(4):554–565.

[39] May A. Structural Brain Imaging: A Window into Chronic Pain. The Neuroscientist 2011;17(2):209–220.

[40] Miller ME, Hui SL, Tierney WM. Validation techniques for logistic regression models. Statistics in medicine 1991;10(8):1213–1226.

[41] O’Reilly RC. The What and How of prefrontal cortical organization. Trends in neurosciences 2010;33(8):355–361.

[42] Olesen SS, Brock C, Krarup AL, Funch-Jensen P, Arendt-Nielsen L, Wilder-Smith OH, Drewes AM. Descending inhibitory pain modulation is impaired in patients with chronic pancreatitis. Clin Gastroenterol Hepatol 2010;8(8):724–730.

[43] Power JD, Barnes KA, Snyder AZ, Schlaggar BL, Petersen SE. Spurious but systematic correlations in functional connectivity MRI networks arise from subject motion. NeuroImage 2012;59(3):2142–2154.

[44] Rajapakse JC, Giedd JN, Rapoport JL. Statistical approach to segmentation of single-channel cerebral MR images. IEEE transactions on medical imaging 1997;16(2):176–186.

[45] Reddan MC, Wager TD. Modeling Pain Using fMRI: From Regions to Biomarkers. Neuroscience Bulletin 2017.

[46] Riederer F, Gantenbein AR, Marti M, Luechinger R, Kollias S, Sándor PS. Decrease of gray matter volume in the midbrain is associated with treatment response in medication-overuse headache: possible influence of orbitofrontal cortex. J Neurosci 2013;33(39):15343–15349.

[47] Robinson M, Boissoneault J, Sevel L, Letzen J, Staud R. The Effect of Base Rate on the Predictive Value of Brain Biomarkers. The journal of pain: official journal of the American Pain Society 2016;17(6):637–641.

[48] Rodriguez-Raecke R, Niemeier A, Ihle K, Ruether W, May A. Brain gray matter decrease in chronic pain is the consequence and not the cause of pain. J Neurosci 2009;29(44):13746–13750.

[49] Sakai K, Rowe JB, Passingham RE. Active maintenance in prefrontal area 46 creates distractor-resistant memory. Nature neuroscience 2002;5(5):479–484.

[50] Satterthwaite TD, Wolf DH, Loughead J, Ruparel K, Elliott MA, Hakonarson H, Gur RC, Gur RE. Impact of in-scanner head motion on multiple measures of functional connectivity: relevance for studies of neurodevelopment in youth. NeuroImage 2012;60(1):623–632.

[51] Seminowicz DA, Moayedi M. The Dorsolateral Prefrontal Cortex in Acute and Chronic Pain. The journal of pain: official journal of the American Pain Society 2017;18(9):1027–1035.

[52] Simons LE, Elman I, Borsook D. Psychological processing in chronic pain: a neural systems approach. Neuroscience and biobehavioral reviews 2014;39:61–78.

[53] Simons LE, Pielech M, Erpelding N, Linnman C, Moulton E, Sava S, Lebel A, Serrano P, Sethna N, Berde C, Becerra L, Borsook D. The responsive amygdala: treatment-induced alterations in functional connectivity in pediatric complex regional pain syndrome. Pain 2014;155(9):1727–1742.

[54] Simons LE, Sieberg CB, Carpino E, Logan D, Berde C. The Fear of Pain Questionnaire (FOPQ): assessment of pain-related fear among children and adolescents with chronic pain. J Pain 2011;12(6):677–686.

[55] Simons LE, Sieberg CB, Conroy C, Randall ET, Shulman J, Borsook D, Berde C, Sethna NF, Logan DE. Children With Chronic Pain: Response Trajectories After Intensive Pain Rehabilitation Treatment. J Pain 2017.

[56] Simons LE, Sieberg CB, Pielech M, Conroy C, Logan DE. What does it take? Comparing intensive rehabilitation to outpatient treatment for children with significant pain-related disability. J Pediatr Psychol 2013;38(2):213–223.

[57] Smucker MR, Craighead WE, Craighead LW, Green BJ. Normative and reliability data for the Children’s Depression Inventory. Journal of abnormal child psychology 1986;14(1):25–39.

[58] Tétreault P, Mansour A, Vachon-Presseau E, Schnitzer TJ, Apkarian AV, Baliki MN. Brain Connectivity Predicts Placebo Response across Chronic Pain Clinical Trials. PLoS Biol, Vol. 14, 2016. p. e1002570.

[59] Tohka J, Zijdenbos A, Evans A. Fast and robust parameter estimation for statistical partial volume models in brain MRI. NeuroImage 2004;23(1):84–97.

[60] Van Dijk KR, Sabuncu MR, Buckner RL. The influence of head motion on intrinsic functional connectivity MRI. NeuroImage 2012;59(1):431–438.

[61] van Velzen GAJ, Rombouts SARB, van Buchem MA, Marinus J, van Hilten JJ. Is the brain of complex regional pain syndrome patients truly different? European Journal of Pain 2016;20(10):1622–1633.

[62] Wager TD, Rilling JK, Smith EE, Sokolik A, Casey KL, Davidson RJ, Kosslyn SM, Rose RM, Cohen JD. Placebo-induced changes in FMRI in the anticipation and experience of pain. Science (New York, NY) 2004;303(5661):1162–1167.

[63] Walker LS, Greene JW. The functional disability inventory: measuring a neglected dimension of child health status. Journal of pediatric psychology 1991;16(1):39–58.

[64] Whitfield-Gabrieli S, Nieto-Castanon A. Conn: a functional connectivity toolbox for correlated and anticorrelated brain networks. Brain connectivity 2012;2(3):125–141.

[65] Wilke M, Holland SK, Altaye M, Gaser C. Template-O-Matic: a toolbox for creating customized pediatric templates. NeuroImage 2008;41(3):903–913.

[66] Woo CW, Krishnan A, Wager TD. Cluster-extent based thresholding in fMRI analyses: pitfalls and recommendations. NeuroImage 2014;91:412–419.

[67] Xu Q-S, Liang Y-Z, Du Y-P. Monte Carlo cross-validation for selecting a model and estimating the prediction error in multivariate calibration. Journal of Chemometrics 2004;18(2):112–120.

[68] Youssef AM, Macefield VG, Henderson LA. Cortical influences on brainstem circuitry responsible for conditioned pain modulation in humans. Hum Brain Mapp 2016;37(7):2630–2644.

